# Contribution of the delta-class glutathione S-transferase to agrochemical adaptation in *Apis mellifera*

**DOI:** 10.1101/2023.07.20.549922

**Authors:** Timothy W. Moural, Sonu Koirala B K, Gaurab Bhattarai, Ziming He, Haoyang Guo, Ngoc T. Phan, Edwin G. Rajotte, David J. Biddinger, Kelli Hoover, Fang Zhu

## Abstract

The European honey bee, *Apis mellifera*, serves as the principle managed pollinator species globally. In recent decades, honey bee populations have been facing serious health threats from combined biotic and abiotic stressors, including diseases, limited nutrition, and agrochemical exposure. Understanding the molecular mechanisms underlying xenobiotic adaptation of *A. mellifera* is critical, considering its extensive exposure to phytochemicals and agrochemicals present in flowers, propolis, hives, and the environment. In this study, we conducted a comprehensive structural and functional characterization of AmGSTD1, a delta class glutathione S-transferase (GST) enzyme, to unravel its roles in agrochemical detoxification and antioxidative stress responses. Significantly, we determined the 3D structure of a honey bee GST using protein crystallography for the first time, providing new insights into its molecular structure. Our investigations revealed that AmGSTD1 efficiently metabolizes model substrates, including 1-chloro-2,4-dinitrobenzene (CDNB), p-nitrophenyl acetate (PNA), phenylethyl isothiocyanate (PEITC), propyl isothiocyanate (PITC), and the oxidation byproduct 4-hydroxynonenal (4-HNE). Moreover, we discovered that AmGSTD1 exhibits binding affinity with the fluorophore 8-Anilinonaphthalene-1-sulfonic acid (ANS), which can be inhibited with various herbicides, fungicides, insecticides, and their metabolites. These findings highlight the potential contribution of AmGSTD1 in safeguarding honey bee health against various agrochemicals and their metabolites, while also mitigating oxidative stress resulting from exposure to these substances.

## 1. Introduction

The honey bee (*Apis mellifera*), known for its honey production and vital role in pollinating a wide range of plants, including both cultivated crops and wild flora, is of paramount importance to the well-being of human societies and ecosystems [1–5]. Unfortunately, in recent decades, honey bee health has been significantly challenged by various environmental stresses [6, 7]. These factors might include stresses encompassing both biotic factors such as pathogens and parasites, as well as abiotic factors including xenobiotics, undernourishment, climate change, and heavy metals, among others [7]. As generalist pollinators, honey bees have the potential to be exposed to a broad diversity of phytochemicals. Exposure occurs when they collect floral rewards such as nectar and pollen or when they encounter antimicrobial plant resins [8, 9]. In agroecosystems, honey bees are also likely to be exposed to numerous agrochemicals, including insecticides, herbicides, and fungicides [10, 11]. Accumulated studies have provided significant insights into the acute toxicity and sublethal impacts of xenobiotic exposure on bees’ foraging, learning, memory, gut microbiota, and immunity [12–23]. Interestingly, when present at low concentrations, certain xenobiotics have been found to exhibit hormetic effects that can benefit honey bee health [24–26]. Therefore, gaining a thorough understanding of the mechanisms underlying honey bees’ adaptation to xenobiotics becomes crucial.

Arthropods adapt to xenobiotics through different mechanisms such as a thickened cuticle [27–29], behavioral resistance [30–32], target site insensitivity [33, 34], and enhanced metabolic detoxification [35–39]. Compared with the genomes of the fruit fly *Drosophila melanogaster* and the African malaria mosquito *Anopheles gambiae*, the honey bee genome has a smaller number of cuticular protein-encoding genes responsible for developing complex cuticles [40]. It is hypothesized that such reduction might be due to their settlement in a protective colony. In terms of behavior, three outcomes can be observed: a) avoidance of contaminated food (negative effect) reducing exposure to harmful substances [19, 41], b) no change in behavior (no effect) [42], and c) attraction to contaminated food (positive effect) [43]. The latter two outcomes result in exposure to toxic compounds, which can pose a risk to bees’ health. Target site insensitivity mediated xenobiotic adaptation has been extensively reported in arthropods such as mosquito [44–46], bed bug [47–49], peach potato aphid [50, 51], cotton bollworm [52], Colorado potato beetle [53, 54], two-spotted spider mite [38, 55, 56], monarch butterfly [57–59], and honey bee [60]. This mechanism reduces or eliminates the binding affinity of chemicals to their target proteins and plays a critical role in xenobiotic adaptation. Arthropods also defend against xenobiotic compounds through enhanced metabolic detoxification [35, 36]. The known enzymes associated with metabolic detoxification or elimination of xenobiotics are the cytochrome P450 monooxygenases (P450s), glutathione S-transferases (GSTs), carboxylesterases (COEs), ATP-binding cassette (ABC) transporters, and UDP-glycosyltransferases (UGTs) [61–65].

One of the major detoxification enzymes recently studied in the honey bee is cytochrome P450 monooxygenases (P450s) [66–68]. P450s constitute a superfamily of heme-thiolate enzymes that participate in both biosynthesis of endogenous compounds and xenobiotic detoxification in insects [67, 69–71]. For example, CYP9Q2 and CYP9Q3 was recently demonstrated to play a key role in detoxifying N-cyanoamidine neonicotinoids [67]. In addition, honey bee diets (honey, bee bread, nectar and pollen) contain various phytochemicals such as p-coumaric acid and quercetin, which induce the expression of CYP6AS and CYP9Q P450s and protect honey bees against pathogens and pesticides [66, 72]. Another important group of detoxification enzymes involved in detoxifying endo- and xenobiotic compounds are GSTs. GSTs make up a superfamily of multifunctional proteins mainly catalyzing the reaction of nucleophilic attack of the thiol group of reduced glutathione (GSH) to electrophilic substrates or toxic chemical compounds previously processed in the phase I detoxification process carried out by P450s and/or COEs [64, 65, 73]. GSTs also have peroxidase activity, which reduces oxidative stress and sequestrates xenobiotics through substrate binding [63].

GSTs are classified based on their cellular location: mitochondrial GSTs, microsomal GSTs, and cytosolic GSTs [73–76]. The largest family, the cytosolic GSTs, is divided into six subclasses: delta, epsilon, omega, sigma, theta, and zeta [75, 77]. Among them, epsilon and delta classes are insect-specific and are the major GSTs involved in xenobiotic detoxification [78]. The honey bee genome contains only 10 putatively functional GST genes and has only one delta class GST (AmGSTD1) and no epsilon GSTs [61]. It was hypothesized that the scarcity of delta and epsilon GSTs in honey bees make them more sensitive to toxic chemical compounds [61]. However, a recent study suggested that honey bees were not more sensitive than other insects to the range of insecticides used in the environment despite their smaller number of detoxification enzymes [79]. In a transcriptome study, AmGSTD1 was upregulated by exposure to pesticides that were frequently used in beehives, suggesting it may play an important role in pesticide detoxification [80]. However, the functional and structural characterization of AmGSTD1 is still lacking.

This study aimed to solve the three-dimensional structure of AmGSTD1 through x-ray crystallography. We have determined the crystal structure of AmGSTD1 complexed with GSH at 1.54 Å resolution. Site directed mutagenesis was used to determine specific amino acids that facilitate binding of GSH with hydrophobic substrates in the “G-site” and “H-site”. We also performed disc diffusion assays to determine the antioxidative properties of this protein and investigated the gene expression patterns of AmGSTD1 in nurse and forager bees. In addition, the binding affinity of AmGSTD1 to 25 commonly used phytochemicals and agrochemicals was determined using a fluorescence competitive binding assay. This study provides a thorough understanding of the structure and function of the only delta GST protein characterized to date in honey bee and demonstrates that AmGSTD1 potentially plays an essential role in protecting honey bee health from the stresses of various agrochemicals.

## 2. Materials and methods

### 2.1. Recombinant AmGSTD1 protein expression and purification

The full-length *AmGSTD1* coding sequence (XP_006563394.1) was introduced into the pET9Bc plasmid vector with ligation independent cloning and then the constructed plasmid was used to transform the Rosetta^TM^ II (DE3) pLysS BL21 *E. coli* expression strain. The pET9Bc expression vector was obtained from the Addgene nonprofit plasmid repository and was gifted to the repository by Scott Gradia (Addgene plasmid # 48285; http://n2t.net/addgene:48285; RRID: Addgene_48285). For protein expression, bacterial cells containing pET9Bc-*AmGSTD1* were incubated at 37 °C overnight at 250 rpm in 100 ml TB media (2 g of tryptone, 2.4 g of yeast extract, 0.4 mL of glycerol, 10 mL of 10x PBS, and 90 ml of diH_2_O). The overnight culture was used to inoculate 2 L of TB media and then incubated at 37 °C until its optical density (OD_600nm_) reached 0.4-0.6. Next, protein expression was induced by the addition of IPTG (0.5 mM) and the culture was further incubated in a shaking incubator set at 250 rpm and 20 °C for a further 20-24 h. Bacterial cells were then harvested by centrifuging (4000 rpm) at 4 °C, then the supernatant was removed, and the pellet was stored at -20°C for later use. For purification, cells were lysed by sonication (Branson Digital Sonifier SFX 150) after resuspension in buffer containing 20 mM NaPi, 500 mM NaCl, 3.3 mM NaN_3,_ 20 mM Imidazole, pH of 7.6, and a Pierce Protease Inhibitor Tablet (Thermo Scientific™). The lysate was then centrifuged at 18000 rcf and the supernatant was split off and injected onto a Co-NTA column attached to a Bio-Rad NGC chromatography system (Bio-Rad Laboratories, Hercules, CA, USA). The Co-NTA column was washed with lysis buffer until a 280 nm reading of the flowthrough returned to baseline. Then the AmGSTD1 protein was eluted with buffer containing 20 mM NaPi, 300 mM NaCl, and 250 mM Imidazole (pH 7.6). The elution was then 100 × fold buffer exchanged with 5 mM NaPi, 5 mM HEPES, pH 6.8 and a centrifugal concentrator. Next, the concentrated AmGSTD1 protein was injected onto a Hydroxyapatite column (HA) connected to the NGC chromatography System (Bio-Rad Laboratories, Hercules, CA, USA). A gradient of 5 mM NaPi and 5 mM HEPES, pH 6.8 to 500 mM NaPi (pH 6.8) was used to wash and elute AmGSTD1. Next, the AmGSTD1 protein fractions were combined, and the buffer exchanged into 20 mM Tris, 150 mM NaCl, 1 mM DTT, 1 mM EDTA (pH 7.6), then injected onto a Protein Ark HiFliQ GST FPLC column connected to the NGC chromatography system. The column was washed until the 280 nm reading returned to baseline and then eluted with 20 mM Tris, 150 mM NaCl, 1 mM DTT, 1mM EDTA, 20 mM reduced glutathione (pH 7.6). For experiments requiring apo protein, the GST column was skipped and chromatography with a Cytiva® HiPrep Sephacryl S-200 HR size exclusion column with 20 mM HEPEs and 150 mM NaCl (pH 7.6) was used as the final purification step. Chromatography fractions were analyzed by sodium dodecyl sulfate polyacrylamide gel electrophoresis (SDS-PAGE) after each step of purification to examine the quality and quantity of protein. Final purified protein samples exhibited a single band when analyzed by SDS-PAGE. The protein concentration of the purified protein was measured on a NanoDrop™ One (Invitrogen™) using an absorption coefficient of 1.184 L g-1 cm-1 at λ_280nm_.

### 2.2. X-ray crystallography

AmGSTD1 crystals were grown by sitting drop vapor diffusion at 18°C. Highly purified AmGSTD1 protein at a concentration of 20mg/ml, in a buffer composed of 20 mM HEPES, 150 mM NaCl, 10mM GSH, and 2mM DTT (pH of 7.2) was mixed with reservoir solution at the ratio of 1:1 in a sitting drop well. The mixture was incubated against mother liquor reservoir solution (100 mM MES pH 6.5, 100mM NaCl, 25% PEG 4K). AmGSTD1 crystal diffraction data were collected at the 19-ID beamline of Argonne National Laboratory’s Advanced Photon Source Structural Biology Center with a Pilatus 6M detector. The CCP4 software package with DIALS, xia2 software was used for processing diffraction data [81–85]. Phasing was done by using nlGSTD (PDB: 3WYW) as a search model with Phaser implemented in Phenix [86–88]. Refinement and model building were performed by using Phenix and Coot [88, 89]. Search models for molecular replacement were identified by a NCBI blastp with the AmGSTD1 amino acid sequence search against the Protein Data Bank (PDB) wwpd.org [90–94]. Structural analysis and figures of AmGSTD1 were created using UCSF Chimera, UCSF Chimera X, and Coot [89, 95–97]. Secondary structure was analyzed by running the DSSP command in ChimeraX that uses the Defining the Secondary Structure of Proteins algorithm [96, 98]. The amino acid sequence was numbered according to the first methionine of the AmGSTD1 sequence. The coordinates and structure factors for the final model of AmGSTD1 complexed with GSH was deposited in the PDB under accession code 7RHP.

### 2.3. Enzymatic assay

Enzyme kinetics of purified recombinant AmGSTD1 was studied using reduced glutathione (GSH), 1-chloro-2,4-dinitrobenzene (CDNB), p-nitrophenyl acetate (PNA), phenylethyl isothiocyanate (PEITC), propyl isothiocyanate (PITC), and 4-hydroxynonenal (4-HNE). For the GSH assay, the concentration of GSH was varied from 0.25 mM to 3 mM while holding the CDNB concentration constant at 1 mM. For the other substrates concentrations ranged from 0.25 mM to 3 mM for CDNB, from 0.25 mM to 3 mM for PNA, from 0.25 mM to 2 mM for PEITC, and from 0.25 mM to 4 mM for PITC, while holding GSH concentration constant at 4 mM. For the assay with 4-HNE, the concentration of GSH was held constant at 0.5 mM, and 4-HNE was varied from 0.0234 mM to 0.375 mM. The concentration of AmGSTD1 was 0.05 mg/mL for the CDNB and GSH assay, 0.025 mg/mL for the PNA and 4-HNE assays, and 0.01 mg/mL for the PIETC and PITC assays. Reactions were carried out in 96-well UVstar® microplates (Greiner Bio-One). The reaction buffer was 100 mM KPi buffer, pH 6.5. Assays with CDNB, GSH, PNA, PEITC, PITC, and 4-HNE were run in reaction volumes of 200 μL in triplicate. Additionally, control reactions minus AmGSTD1 were run to account for non-enzymatic reaction contributions. The reactions were monitored by measuring change in absorbance for 1 minute continuously, with 10 second reads at a constant temperature of 30°C, using a Tecan Spark® multi-mode plate reader. The kinetic parameters and Michaelis–Menten plots were analyzed and generated using GraphPad Prism 9.5.1. The wavelengths used and conjugated product concentrations for CDNB, PNA, PIETC, and PITC or substrate depletion reactions for HNE were calculated by path-length corrected molar attenuation coefficients published previously [99–102].

### 2.4. Site directed mutagenesis

A total of 8 mutants targeted at the H-site and one mutant targeted at the G-site were created by substituting key amino acid residues in the AmGSTD1 active site identified from the protein crystal structure. The PCR primers (Table S1) were designed using NEBaseChanger to generate substituted nucleotides in the target sequence of *AmGSTD1*. PCR-based mutagenesis was carried out using the Q5^®^ Site-Directed Mutagenesis Kit (New England Biolabs^®^, MA, USA) as per the manufacturer’s recommendations. The wildtype AmGSTD1 was used as template and the resulting site-directed mutants were sequenced to verify the correct mutations. The mutant pET9Bc-AmGSTD1 constructs were then transformed into Rosetta^TM^ II (DE3) pLysS cells for protein expression. The mutant AmGSTD1 proteins were expressed and purified in the same manner as the wildtype AmGSTD1. After purification, the mutant AmGSTD1 proteins were submitted to enzyme kinetics assays with GSH and CDNB as described in section 2.3.

### 2.5. Fluorescence binding assay

8-Anilinonaphthalene-1-sulfonic acid (ANS) was used as a fluorescent probe to assess ligand binding to the H-site of AmGSTD1. ANS has been previously used in displacement based competitive fluorescence binding assays to investigate the binding of ligands to hydrophobic pockets within proteins [103]. For GSTs, ANS has been shown to bind the proteins in a 1:1 molar ratio and has specificity for the H-site while not disturbing the proteins conformational stability [104, 105]. A saturation binding assay of AmGSTD1 with ANS was performed at a constant concentration of protein at 1.7 μM and the concentration of ANS was varied from 0 μM to 74.25 μM. For ANS displacement studies, AmGSTD1 was held at 1.7 μM, ANS at 50 μM, and the competitor ligands varied from 0 μM to 750 μM, with upper limits being lower for some ligands due to solubility issues. All assays were run in 96-well flat bottom black plates with a total volume of 200 μL binding reaction. A multi-mode Tecan Spark® plate reader and fluorescence intensity were measured using a 380nm/20 nm excitation filter and 485 nm/20nm emission filter. Competitor ligands included the known GST substrates GSH (negative control for displacing ANS form the H-site), CDNB and ethacrynic acid (positive controls for displacing ANS from the H-site). Herbicides (paraquat, dicamba, atrazine, triclopyr, the metabolite of triclopyr 3,5,6-trichloro-2-pyridinol (TCP), and 2,4-D), fungicides (chlorothalonil, propiconazole, and myclobutanil), neonicotinoids (dinotefuran, imidacloprid and metabolite 6-chloronicotinic acid), pyrethroids (permethrin, fenvalerate, deltamethrin) and the metabolite 3-phenoxybenzaldehyde, organophosphates (malathion, coumaphos, chlorpyrifos, and diazinon), a carbamate (carbaryl), and an acaricide (amitraz), along with natural products (p-coumaric acid, caffeine, and nicotine) were used as ligands for the binding reactions. GraphPad Prism 9.5.1 was used for curve fitting the saturation binding curve and calculation of the AmGSTD1/ANS *K_d_*. Fluorescence inhibition curves were generated to calculate IC_50_ values for competitive ligands and used to calculate dissociation constants (K_i_) with the equation *K_i_* = [IC_50_]/(1 + [ANS]/*K_d_*-ANS) [106]. The assays were run in triplicate.

### 2.6. Disc diffusion assay

Disc diffusion assays were performed following Burmeister’s protocol [107]. The pET-9BC vector only (i.e., control group) and recombinant pET-9BC-AmGSTD1 (treatment group) expressed in Rosetta^TM^ II (DE3) pLysS were cultured in LB medium at 37 °C and 250 rpm. The control and the treatment group were grown and expressed separately after adding 1 mM of IPTG when the OD_600_ reached 0.6. Then, the cultures were expressed by incubating at 37 °C for 7 hours. Approximately 5 × 10^8^ cells were plated on each LB agar plate with the antibiotics, ampicillin (200 mg/ml) and chloramphenicol (30 microgram/ml). Then the plates were incubated at 37 °C for 1 hour. After that, filter discs (6-mm diameter) soaked with different concentrations of cumene hydroperoxide or hydrogen peroxide (0 mM, 50 mM, 100 mM, 150 mM and 200 mM) and different concentrations of paraquat (0 mM, 50 mM, 200 mM, 400 mM and 600 mM) were placed on the plates separately. Lastly, the halo diameter from the control and treatment plates was recorded after 12 hours of incubation at 37°C.

### 2.7. RNA extraction, cDNA synthesis, and qRT-PCR

The honey bees were sourced from three different hives located 60 m apart from each other in the Penn State Wiley Apiary (University Park, PA). These hives have been maintained based on best management practices (https://extension.psu.edu/best-management-practices-for-bee-health; accessed on July 20^th^, 2023). The nurse bees were selected by shaking frames of brood, and the forager bees were selected by shaking honey frames. Subsequently, all bees were promptly collected and immediately frozen using liquid nitrogen. The bee samples were then stored at -80°C till use. The nurse or forager bees were dissected in phosphate-buffered saline solution. Different tissues such as heads, fat bodies, Malpighian tubules, midguts, legs, and muscles were harvested for the experiment. The total RNA of nurse or forager honey bee tissue samples was extracted using the Invitrogen TRIzol reagent (Invitrogen, Carlsbad, CA, USA). RNA quantity and quality were determined by a NanoDrop 2000 spectrophotometer (NanoDrop, Wilmington, DE, USA). The A260/280 value of 1.8-2.0 was used as a standard to assess RNA quality. Then, the complementary cDNA was synthesized with an M-MLV Reverse transcriptase (Thermo Fisher Scientific, Waltham, MA). The synthesized cDNA was used as a template for qRT-PCR reactions. PCR conditions included 3 min at 95°C followed by 39 cycles of 10 s at 95°C, and 55°C for 30 s. qRT-PCR was conducted with 1 μL of cDNA, 5 μL of FastStart SYBR Green Master (Roche Diagnostics, Indianapolis, IN USA), 0.4 μL of qRT-PCR primers (Table S1), and 3.6 μL of ddH_2_O in a 10 μL total reaction volume using Bio-Rad CFX Connect™ Real-Time PCR System (Bio-Rad, CA, USA). The expression of the housekeeping gene GADPH (XM393605) was used for normalization (Table S1) [108]. The relative *AmGSTD1* gene expression in different tissues was calculated in reference to the tissue with the lowest expression level by the 2^-ΔΔCT^ method [109]. Each reaction performed had three biological and two technical replications. The overall difference in the level of expression among tissues was analyzed by one-way ANOVA with Tukey HSD test for multiple comparisons in R (Version 4.1.0).

### 2.8. AmGSTD1 cloning and phylogenetic analysis

Specific forward and reverse primers (Table S1) were designed to amplify the full length of the *AmGSTD1* coding sequence. The potential gene was amplified using the cDNA obtained in the previous steps. The PCR products obtained were purified using DNA clean and Concentrator^TM^-25 (Zymo Research, CA, USA). The purified products were further treated with T4 DNA polymerase. Likewise, the pET-9BC vector (Addgene plasmid #48285) digested with restriction enzyme *HpaI* overnight was treated with T4 DNA polymerase. The T4 treated *AmGSTD1* gene was ligated into a T4 treated linearized vector at 2: 8 (vector: PCR product) ratio. Then the reaction was transformed into DH5-alpha competent cells to replicate the newly constructed plasmid and grown on LB (tryptone: 5 g, NaCl: 5 g, Yeast extract: 2.5 g, bactoagar: 7.5 g in 500 ml of diH_2_O) agar plates with selected antibiotics (ampicillin: 200 mg/ml) overnight (16-18 h). Positive colonies were verified using T7 primers (Table S1), then cultured in liquid LB media (tryptone: 5 g, NaCl: 5 g, yeast extract: 2.5 g in 500 ml of diH_2_O) with ampicillin (200 mg/ml) overnight at 37 ℃. The plasmids were extracted and sequenced by Functional Biosciences, Inc. To investigate the phylogenetic relationships of AmGSTD1 with other insect GSTs, a phylogenetic tree was constructed. 182 GST amino acid sequences (characterized and predicted) from five different insect species (*Ap. melifera, Bombyx mori*, *D. melanogaster*, *An. gambiae*, and *Tribolium castaneum*) were retrieved from the NCBI database (Table S2). Multiple sequence alignment was performed using ClustalW [110] in MEGA X [111] with default parameters (gap open penalty: 10, gap extension penalty: 0.2). The maximum likelihood unrooted phylogenetic tree was inferred using RaxML 8.2.12 [112]. Bootstrap analysis was performed in 500 replicates to infer the consensus tree.

## 3. Results

### 3.1. X-ray crystal structure of AmGSTD1

A NCBI blastp search with the AmGSTD1 sequence revealed that the highest identity match was *Nilaparvata lugens* nlGSTD (PDB 3WYW) with a sequence identity of 58.85%, which was subsequently used for molecular replacement [90, 93]. AmGSTD1 crystalized in space group P6_5_22, unit cell a = 60.09 (Å), b = 60.09 (Å), c = 232.05 (Å), α = 90°, β = 90°, γ = 120°. Diffraction data up to 1.54 (Å) was used for structural refinement with R_work_ of 19.03% and R_free_ of 20.23% (Table 1). There was one molecule of AmGSTD1 complexed with GSH per asymmetric unit. Further analyses with PISA revealed the probable biological assembly to be a dimer [113, 114].

**Table 1.** Data collection and refinement.

| Parameter | AmGSTD1 complex with GSH |
| --- | --- |
| **Data Collection** |  |
| Wavelength (Å) | 0.97918 |
| Resolution range (Å) | 52.04 - 1.54 (1.595-1.54) |
| Space group | P 6<sub>5</sub> 2 2 |
| a, b, c (Å) | 60.090, 60.090, 232.050 |
| Total reflections | 624835 (34777) |
| Unique reflections | 37014 (2976) |
| Multiplicity | 16.9 (11.7) |
| Completeness (%) | 95.00 (73.91) |
| *I*/$\sigma$(*I*) | 14.34 (0.34) |
| R-merge | 0.07798 (0.9732) |
| R-meas | 0.08041 (1.017) |
| R-pim | 0.01925 (0.2854) |
| CC1/2 | 1 (0.772) |
| CC* | 1 (0.933) |
| **Refinement** |  |
| Reflections used for R-free | 1666 (128) |
| R-work | 0.1903 (0.2507) |
| R-free | 0.2023 (0.2904) |
| CC (work) | 0.960 (0.861) |
| CC (free) | 0.962 (0.777) |
| No. of non-hydrogen atoms | 1956 |
| Macromolecules | 3416 |
| Ligands | 20 |
| Solvent | 317 |
| Protein residues | 422 |
| Average B-factor | 27.49 |
| Macromolecules | 26.7 |
| Ligands | 41.50 |
| Solvent | 35.2 |

Parentheses placed around values for highest resolution shell.

The AmGSTD1 monomer consisted of two domains, the N-term domain and the C-term domain connected by a linker region coil (Fig. 1). The N-term domain exhibits a thioredoxin-like fold with four β-strands, three α-helices, and two 3_10_-helices. The N-term domain secondary structure was ordered starting with β1 (residues D27-Q30), followed by α1 (residues P35-A46), β2 (residues N52-Q55), 3_10_-1 (residues L58-N60), 3_10_-2 (residues E62-L64), α2 (residues P66-I71), β3 (residues T79-D82), β4 (residues F85-L87), and α3 (residues S90-Y101). Following the last α-helix of the N-term domain there was a 10 amino acid linker loop consisting of residues G102-D111 connecting to the C-terminal domain. The C-terminal domain was a helical domain, composed of five α -helices interspersed with loops and one 3_10_-helix. Directly following the linker loop region was α4 (residues L112-P142) with a bulge close to the middle of α4 at S128, α5 (residues Q149-F165), α6 (residues L197-A193), α7 (residues K202-H211), 3_10_-3 (L212-G214), and α8 (residues Y218-L232). There was a solvent exposed pocket located between the N-term domain and the C-terminal domain. Previously characterized GSTs have been described as having two binding sites within the pocket between the N and C-terminal domains, which have been termed the “G-site” responsible for binding GSH and the “H-site” that binds hydrophobic substrates [115]. A molecular surface representation color coded by lipophilicity revealed that for AmGSTD1 there is a more hydrophilic region on the N-term domain side of the pocket; in this more hydrophilic region of the solvent exposed pocket, clear electron density was seen representative of a complexed glutathione molecule, clearly establishing this location as the “G-site”. Adjacent to the bound GSH was a more lipophilic region that made up the hydrophobic substrate binding pocket, sometimes termed the “H-site” (Fig. 2B and C).

**Figure 1.**
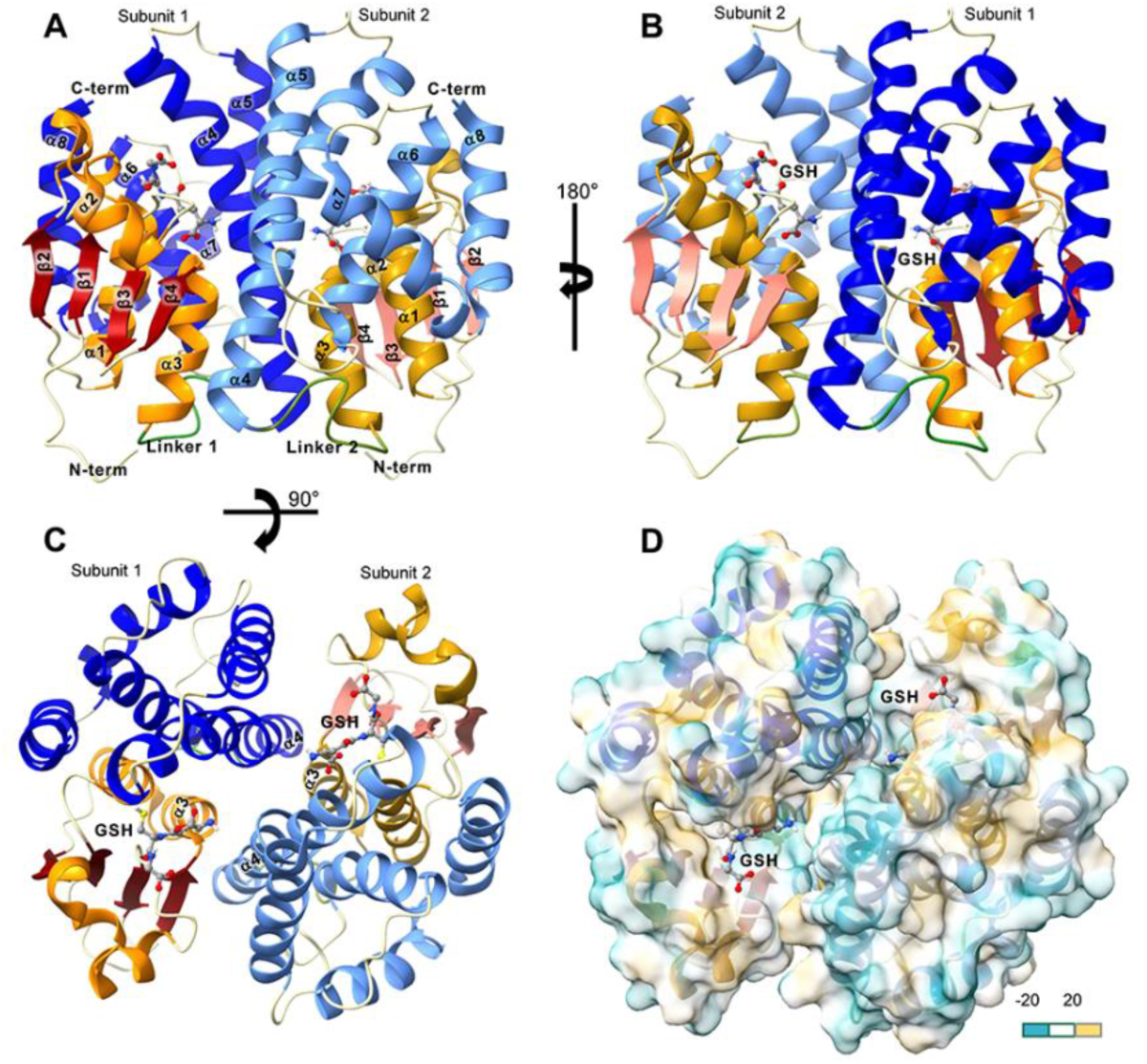
Global structure of AmGST-1 shown as a dimer and represented in ribbon diagram alongside and surface overlayed on a ribbon diagram. (**A**) AmGSTD1 dimeric structure showing Subunit 1 (monomer 1) and Subunit 2 (monomer 2) as a ribbon diagram. For reference, the helices and β-strands have been labeled in order starting from the N-terminus and leading to the C-terminal end. For Subunit 1, the N-terminal domain helices are colored orange, and the β-strands are red, The Linker 1 loop connecting the N-terminal domain to the C-terminal domain is colored green, and the C-terminal helical domain is blue. In Subunit 2 the N-terminal domain helices are colored golden rod and β-strands are salmon, and in the C-terminal domain of Subunit 2 the helices are colored cyan. The crystal complexed glutathione molecule is shown in ball and stick format and colored by element. The same color scheme holds for all the Figure 1 ribbon diagrams. **(B)** The ribbon diagram of the dimeric AmGSTD-1 structure from Figure 1A has been rotated 180°, pulling from left to right about a central axis, to illustrate the 2-fold symmetry axis of the dimer. **(C)** The ribbon diagram of the dimeric AmGSTD-1 structure from Figure 1A has been rotated 90° by pulling down from the top to show the symmetry of the dimer from an additional orientation. **(D)** The ribbon diagram shown in Figure 1C has been overlayed with a semi-transparent lipophilic surface representation of the AmGSTD-1 dimer. The surface color scheme scale is set from -20 to 20, with -20 being the most hydrophilic and colored cyan, and 20 being the most hydrophobic and colored tan. Figures were generated with UCSF Chimera X v:1.5.

**Figure 2.**
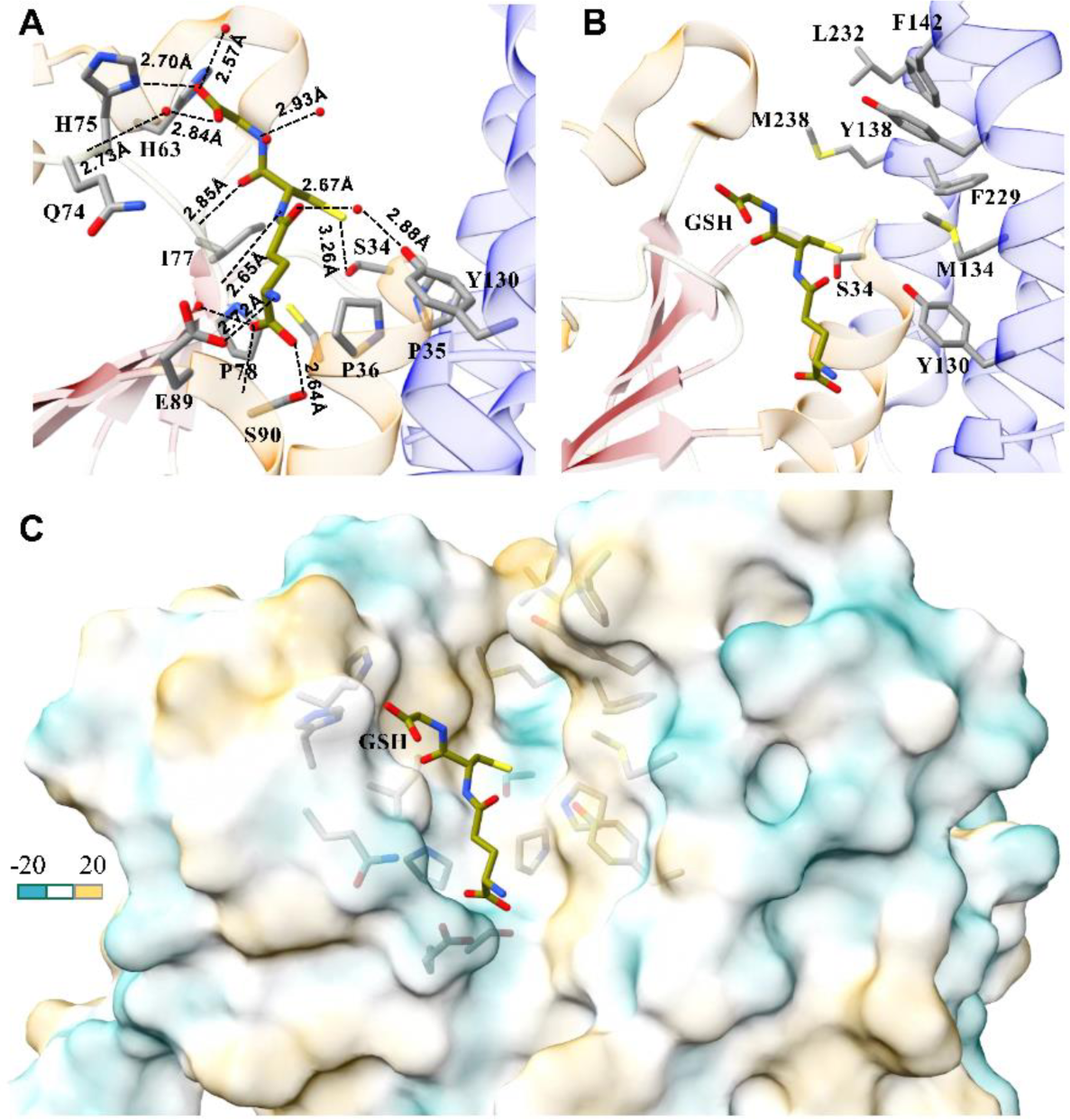
Active site of AmGSTD-1**. (A)** The G-site within the active site of AmGSTD-1 was zoomed in, to illustrate the enzymatically important Ser34 and key ammino acids involved in binding GSH. Sidechains are represented in stick format and colored by element and GSH is colored by heteroatom. Crystallographic water molecules involved in the hydrogen bond network of the G-site with GSH are shown as red spheres. H-bonds are with a dashed line and the respective distance between atoms detailed in the manuscript body are displayed. (**B)** Zoomed in look at the H-site within the active site of AmGSTD-1. Sidechains of residues suspected to be in binding interaction of potential substrates are shown and colored by element. (**C)** Zoomed in lipophilic surface of the AmGSD-1 active site. The complexed GSH molecule is shown and colored by heteroatom. Surface is shown as slightly transparent to allow the important binding interaction sidechains to be seen. The surface color scheme scale is set from -20 to 20, with -20 being the most hydrophilic and colored cyan, and 20 being the most hydrophobic and colored tan. Figures were generated with UCSF Chimera X v:1.5.

### 3.2. Active site of AmGSTD1

The AmGSTD1 active site contains a binding site for glutathione and an adjacent binding site for a hydrophobic substrate. Based on the crystal complex, the glutathione binding site was lined with fifteen amino acids (S34, P36, L58, H63, Q74, H75, T76, I77, P78, E89, S90, R91, M126, C127, Y130) and possessed multiple crystallographic water molecules that participated in an extended hydrogen bonding network with the complexed GSH (Fig. 2A). Key binding interactions between G-site residues and GSH included the sidechain of catalytic Ser34 being located within H-bond distance of the GSH thiol group; specifically, the SER34 OG atom was located 3.26 Å from the SG2 atom of the GSH thiol. Adjacent to the Ser34 hydroxyl was the sidechain thiol of Cys37 (3.46 Å), the Cys37 amide (3.49 Å), the amide of Pro35 (3.14 Å), the amide of Pro36 (3.70 Å), and the Ser34 backbone amide (3.14 Å). In the G-site, residues involved in binding with the γ-glutamyl moiety included Glu89, and Ser90, along with crystallographic waters that formed an extensive hydrogen bond network, including water bridged hydrogen bonds to backbone amides of Ser90 and Arg91. In detail the Glu89 OE1 atom was 2.83 Å from the GSH γ-glutamyl N1 atom, the Ser90 OG atom was 2.64 Å from γ-glutamyl O11, the Ser90 amide N was 2.89 Å from O12 of GSH, and O11 was 3.09 Å from a crystallographic water that bridged hydrogen bonds to the Arg91 amide N (3.01 Å) and Ser90 amide N (3.40 Å). The OE1 γ-carboxyl of GSH was 2.67 Å from a crystallographic water molecule that bridged a hydrogen bond to the phenolic hydroxyl oxygen atom of Tyr130. The glycyl moiety of GSH is bound to AmGSTD1 through a shared hydrogen bonding network that included His75, His63, Gln74, the sidechain of His75 (O32 of GSH 2.7 Å from ND1 of His75), the His75 backbone carbonyl O located 3.0 Å from O32 of GSH, and two water molecules with one located 2.82 Å from O32 and one 2.84 Å from O31. Additional hydrogen bonds between GSH and the H-site included the Ile77 backbone amide N located 2.85 Å from the cysteinyl O2 carbonyl atom of GSH, and the Ile77 backbone carbonyl O positioned 2.74 Å from the cysteinyl N2 atom of GSH N2 by the adjacent cis-Pro78. Adjacent to the G-site was a hydrophobic pocket making up the H-site. Key residues in the H-site, giving it hydrophobic character, included Tyr130, Met134, Tyr138, Phe142, Met228, Phe229, and Leu232 (Fig. 2C).

### 3.3. Biological assembly

The PISA predicted biological dimer assembly based on the AmGSTD1 crystal structure (PDB 7RHP) is stabilized by dimer interface interactions between α3 and α4 of subunit 1 with α3 and α4 of subunit 2. The dimer interface was stabilized by hydrogen bonds between α3 of subunit 1 and α4 subunit 2, which included Gln100 with Lys113, Tyr96 with Lys113, and Thr95 with Asn119, respectively. As shown in the top half of Fig. 3A, the Gln100 OE1 on α3 of subunit 1 and Lys113 NZ on α4 of subunit 2 were positioned at a distance of 3.68Å from each other, and the Tyr96 OH on α3 of subunit 1 and Lys113 NZ on α4 of subunit 2 were positioned at a distance of 3.27Å. Thr95 OG1 on α3 of subunit 1 was located 3.67Å from ND2 of Asn119 on α4 of subunit 2. Equivalent hydrogen bonds were found between α3 of subunit 2 and α4 of subunit 1 and are shown in the bottom half of Fig. 3A. Additionally, the dimer interface was stabilized by hydrophobic interactions similar to what has been previously described in delta class GST enzymes as a “lock-and-key clasp” like interaction [116]. For AmGSTD1 there was an offset pi interaction between the sidechain of Tyr123 from subunit 1 with Tyr123 of subunit 2 forming the “clasp” (Fig. 3B). The Try123 residues were surrounded by Glu89, Arg91, Ala92, Leu122 and Met126, completing the hydrophobic interaction.

**Figure 3.**
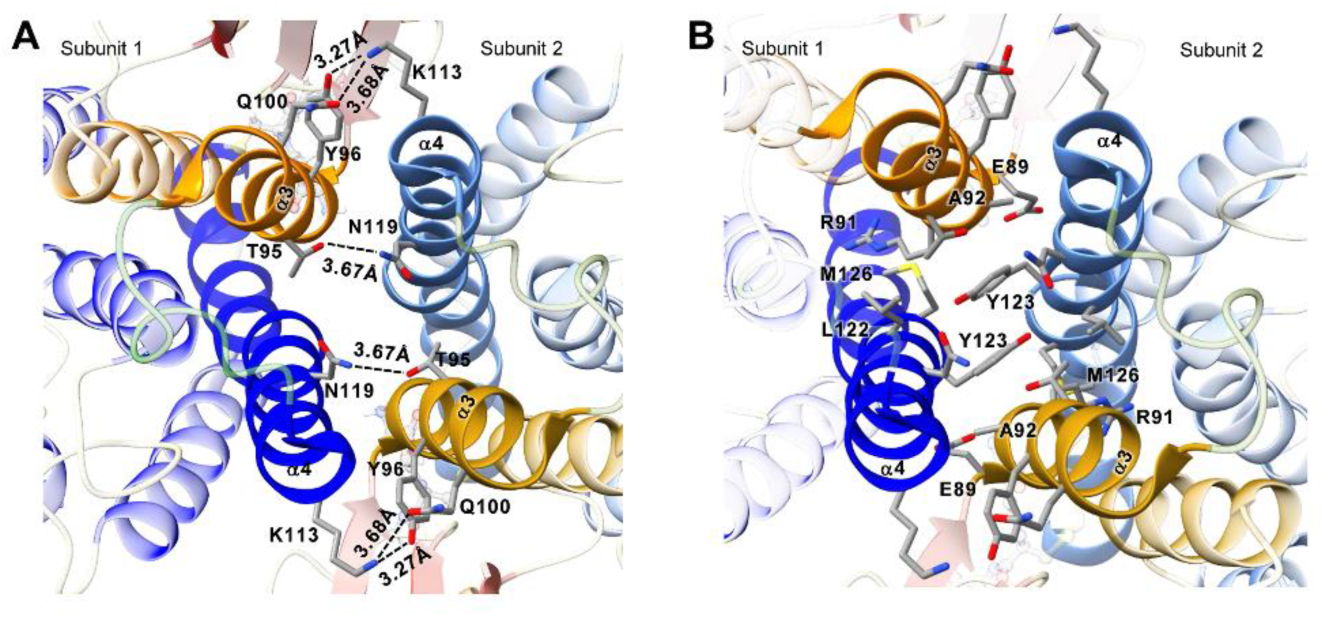
Dimer interface interactions based on PISA analysis of AmGSTD-1 crystal structure data. **(A)** The sidechains of amino acids belong to α3 and α4 are shown and colored by element. The distances between the respective N and O atoms involved in H-bonding interactions between α3 of Subunit 1 and α4 of Subunit 2 are shown, along with the equivalent interactions between α4 of Subunit 1 and α3 of Subunit 2. **(B)** Sidechains of residues involved in hydrophobic interactions at the dimer interface are shown.

### 3.4. Enzymatic and fluorescence binding assays

Kinetic analysis of AmGSTD1 was performed under steady state conditions for CDNB, PNA, PEITC, PITC, HNE and GSH. Kinetic parameters were obtained by nonlinear regression curve fitting to the Michaelis-Menten model with GraphPad Prism 9.5.1. The Michaelis-Menten plots are shown in Fig. 4 and kinetic parameters listed in Table 2. For AmGSTD1, the lowest *K*_m_ was for HNE with a value of 0.106 mM ± 0.034, followed by the model substrate CDNB with a value of 0.525 ± 0.07 mM. *K*_m_ values were 0.819 ± 0.10 mM for PEITC, 1.10 ± 0.14 mM for PITC, 1.16 ± 0.09 mM for PNA, and 1.99 ± 0.19 mM for GSH. The turnover number *k*_cat_ was highest for PITC at 852 ± 39 min^-1^, followed by PNA at 402 ± 14 min^-1^, GSH at 398 ± 20 min^-1^, CDNB at 385 ± 11 min^-1^, and PEITC at 236 ± 12 min^-1^, and the lowest *k*_cat_ was for HNE at 20.7 ± 2.6 min^-1^. Catalytic efficiency (*k*_cat_/*K*_m_) of AmGSTD1 was highest for PITC at 775 min^-1^ mM^-1^, followed by CDNB at 732 min^-1^ mM^-1^, PNA at 345 min^-1^ mM^-1^, PEITC at 288 min^-1^ mM^-1^, GSH at 200 min^-1^ mM^-1^, and HNE at 195 min^-1^ mM^-1^ (Fig. 4; Table 2).

**Figure 4.**
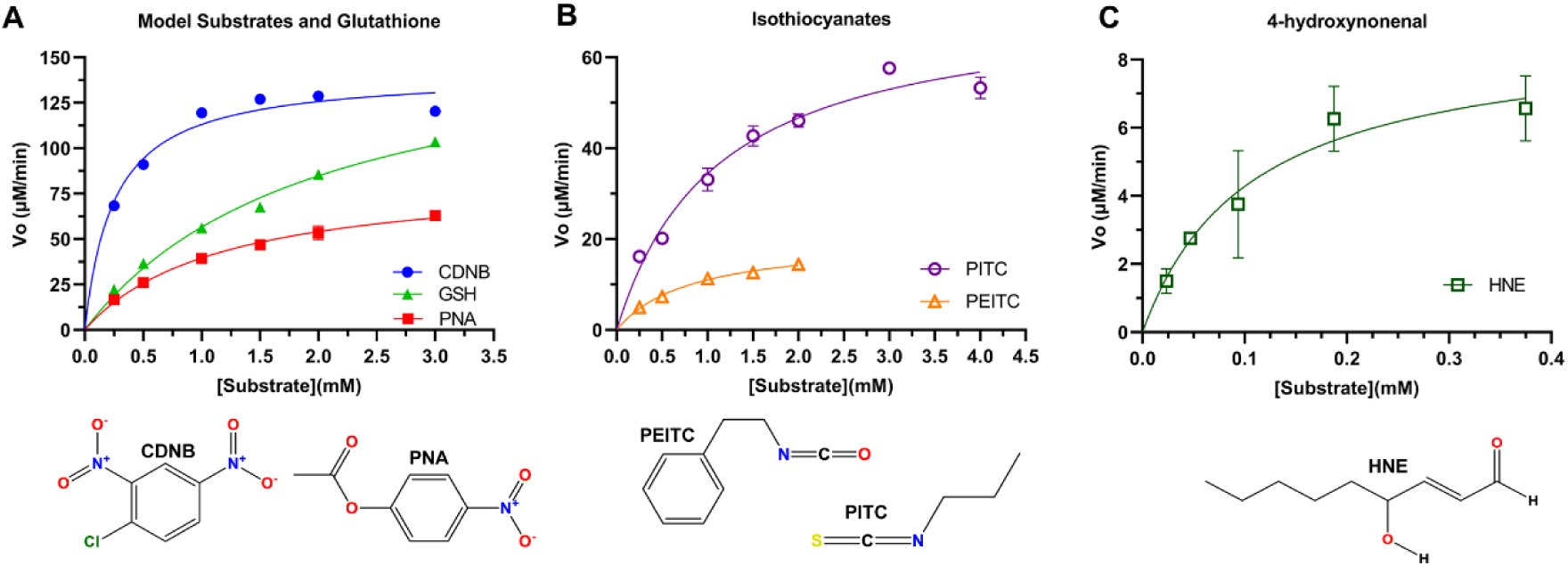
Michaelis-Menten Plots with steady-state initial rates versus substrate concentrations and chemical structures of substrates are shown. (**A)** The conjugation rates of GSH to CDNB and GSH to PNA with concentrations of CDNB and PNA that varied from 0.25 mM to 3 mM, while holding GSH constant at a concentration of 4 mM. For GSH initial velocities, the CDNB concentration was held constant at 1 mM and GSH varied from 0.25 mM to 3 mM. (**B)** The conjugation rates of GSH to PITC (purple circle) and PEITC (orange triangle). GSH was held constant at 4 mM, PITC varied from 0.25 mM to 4 mM, and PIETC varied from 0.25 mM to 2 mM. (**C)** 4-hydroxynonenal varied from 0.0234 mM to 0.375 mM and GSH was held at 0.5 mM to measure conjugation of HNE to GSH. All data were curve fit by non-linear regression analysis and plots generated with GraphPad Prism v:9.5.1 (GraphPad, San Diego, CA, USA).

**Table 2.** Kinetic parameters for wild-type AmGSTD1.

| Substrate | $V_{\max}$ ($\mu\text{M}/\text{min}$) | $K_m$ (mM) | $k_{\text{cat}}$ ($\text{min}^{-1}$) | $k_{\text{cat}}/K_m$ ($\text{min}^{-1} \cdot \text{mM}^{-1}$) |
| --- | --- | --- | --- | --- |
| CDNB | $163 \pm 17$ | $0.525 \pm 0.07$ | $385 \pm 11$ | 732 |
| PNA | $85.5 \pm 2.9$ | $1.16 \pm 0.09$ | $402 \pm 14$ | 345 |
| Phenyl-ITC | $20.1 \pm 0.98$ | $0.819 \pm 0.10$ | $236 \pm 12$ | 288 |
| Propyl-ITC | $72.4 \pm 3.4$ | $1.10 \pm 0.14$ | $852 \pm 39$ | 775 |
| HNE | $8.80 \pm 1.1$ | $0.106 \pm 0.034$ | $20.7 \pm 2.6$ | 195 |
| GSH | $169 \pm 8.4$ | $1.99 \pm 0.19$ | $398 \pm 20$ | 200 |

Site-directed mutagenesis was performed to evaluate active site residues. The mutants created were G-site S34A and H-site Y130A, Y138A, Y138F, F142A, F142Y, M228A, F229Y, and F229E (Table 3). Amongst the alanine active site screening mutants, the G-site S34A mutant exhibited the largest changes in steady state kinetic parameters. When compared to wild type, the S34A *k*_cat_ value decreased by 4.15-fold. The next largest decrease in turnover was for the H-site mutant F142A with 2.08-fold decrease in *k*_cat_ value. Smaller changes in *k*_cat_ were observed for Y130A, F229E, and F229Y, with an increase of 1.15-fold, 1.19-fold, and 1.24-fold, respectively. For Y138F, F142Y, and M228A, *k*_cat_ stayed approximately equivalent with a *k*_cat_ value of the wild-type (Table 3). The wild-type *K_m_* value of 0.525 ± 0.07 mM decreased to 0.480 ± 0.05 mM for Y138A, 0.488 ± 0.03 mM for Y138F, 0.409 ± 0.04 mM for F142A, and to 0.422 ± 0.04 mM for S34A. Increases in *K_m_* were observed for Y130A and F142A to 1.06 ± 0.08 mM, to 1.06 ± 0.07 mM for F229E, to 0.947 ± 0.07 mM for F229Y, and to 0.599 ± 0.06 mM for M228A (Table 3). The only increase in catalytic efficiency (k_cat_/K_m_) was observed for the Y138F mutant. The k_cat_/K_m_ for S34A decreased most (3.33-fold) compared to that of the wild-type (Table 3).

**Table 3.** Kinetic parameters for mutants with the substrate CDNB.

| Mutation | $V_{\max}$ ($\mu\text{M}/\text{min}$) | $K_m$ (mM) | $k_{\text{cat}}$ ($\text{min}^{-1}$) | $k_{\text{cat}}/K_m$ ($\text{min}^{-1} \cdot \text{mM}^{-1}$) |
| --- | --- | --- | --- | --- |
| WT | $163 \pm 17.4$ | $0.525 \pm 0.07$ | $385 \pm 11$ | 732 |
| Y130A | $188 \pm 5.0$ | $1.06 \pm 0.08$ | $443 \pm 12$ | 418 |
| Y138A | $137 \pm 3.5$ | $0.480 \pm 0.05$ | $322 \pm 8.2$ | 669 |
| Y138F | $155 \pm 2.7$ | $0.488 \pm 0.03$ | $365 \pm 6.4$ | 748 |
| F142Y | $163 \pm 5.6$ | $1.06 \pm 0.08$ | $384 \pm 13$ | 629 |
| F142A | $78.7 \pm 1.7$ | $0.409 \pm 0.04$ | $185 \pm 4.1$ | 453 |
| F229E | $194 \pm 4.7$ | $1.06 \pm 0.07$ | $457 \pm 11$ | 433 |
| F229Y | $202 \pm 5.3$ | $0.947 \pm 0.07$ | $476 \pm 11$ | 502 |
| M228A | $156 \pm 4.6$ | $0.599 \pm 0.06$ | $368 \pm 11$ | 614 |
| S34A | $39.5 \pm 0.86$ | $0.422 \pm 0.04$ | $92.8 \pm 2.0$ | 220 |

For the competitive fluorescence binding assay, we first calculated the affinity of the fluorescent probe ANS from the non-linear regression one-site saturation binding curve (Fig. 5), resulting in a calculated *K*_d_ of 2.094 ± 0.102 μM. Next, the inhibition of ANS:AmGSTD1 fluorescence intensity in the presence of competitor ligands was plotted and fit with non-linear regression curves to find the corresponding IC50 values. For GSH, paraquat, dicamba, atrazine, chlorothalonil, propiconazole, dinotefuran, 6-chloronicotinic acid, p-coumaric acid, caffeine, nicotine, carbaryl, amitraz, malathion, coumaphos, diazinon, permethrin, fenvalerate, and deltamethrin, no binding was observed (Fig. 6). For CDNB, EA, triclopyr, 2,4-D, TCP, myclobutanil, imidacloprid, and 3-phenoxybenzadheyde, significant inhibition of fluorescence intensity was observed (Fig. 6). The IC50 values were 1.22 ± 0.055 mM (CDNB), 1.02 ± 0.040 Mm (EA), 1.86 ± 0.19 mM (triclopyr), 1.01 ± 0.056 mM (2,4-D), 0.613 ± 0.027 mM (TCP), 3.64 ± 1.3 mM (myclobutanil), 3.18 ± 0.637 mM (imidacloprid), and 7.40 ± 4.7 mM (3-phenoxybenzadheyde) (Table 4). The IC50 values obtained from the resulting curve fits were then used to find the respective *K*_i_ values using the Cheng and Prusoff equation [117]. The *K*_i_ value was 49.6 μM for CDNB, 41.5 μM for EA, 76.1 μM for triclopyr, 41.4 μM for 2-4-D, 25.0 μM for TCP, 148 μM for myclobutanil, 130 μM for imidacloprid, and 0.604 mM for 3-phenoxybenzaldehyde (Table 4).

**Figure 5.**
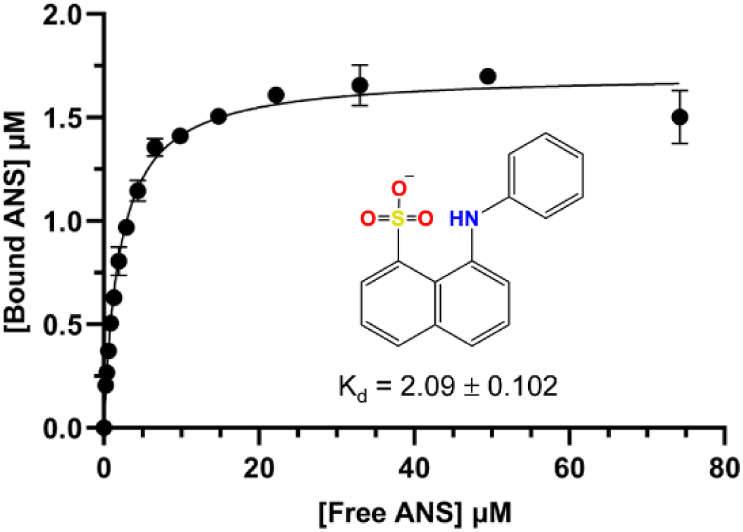
Binding of the fluorescent reporter 8-Anilinonaphthalene-1-sulfonic acid (ANS) to recombinant AmGSTD-1. A solution of AmGSTD-1 protein was titrated with the fluorescent reporter ANS in 96-well flat black microtiter plates. The fluorescence intensity was measured using a 380 nm/20 nm excitation filter and 485 nm/20nm emission filter. The data were analyzed with GraphPad Prism software by using a one-site specific binding model. The chemical structure of the fluorescent reporter ANS is shown, and the calculated K_d_ with SE is reported.

**Figure 6.**
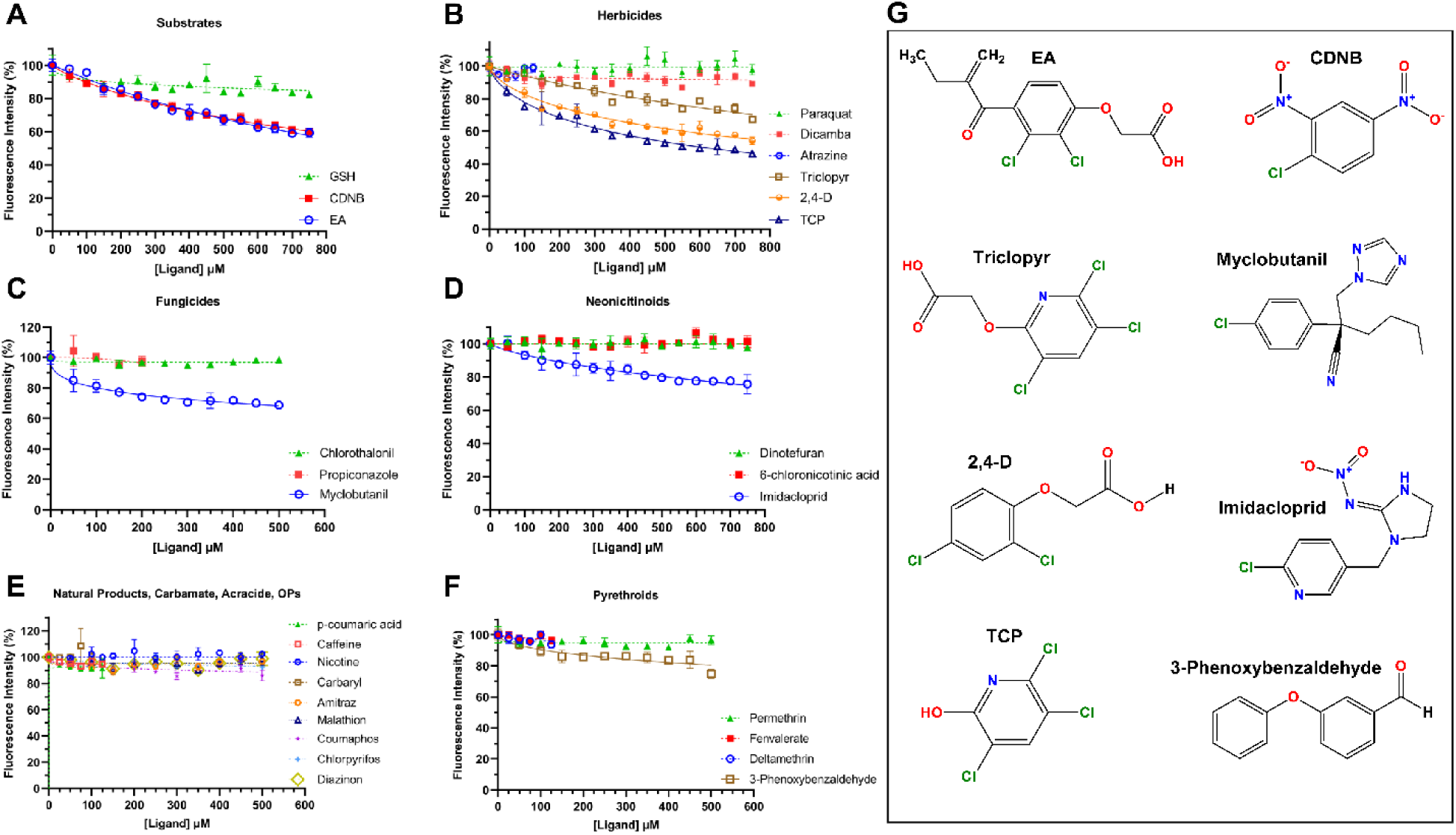
Competitive binding curves of various potential ligands of AmGSTD-1, showing the percentage of fluorescence intensity recorded in the presence of the ligand vs. the concentration of ligand relative to controls with no ligand. AmGSTD-1 (1.7 μM), ANS (50 μM), and ligands were added to 96-well flat black microtiter plates at the final concentrations indicated, and assays were performed in triplicate. **(A)** Competitive binding curve showing fluorescence intensity in the presence of known glutathione S-transferase substrates. **(B)** Competitive binding curve showing fluorescence intensity in the presence of multiple herbicides and the known metabolite TCP. **(C)** Competitive binding curve showing fluorescence intensity in the presence of multiple fungicides. **(D)** Competitive binding curve showing fluorescence intensity in the presence of neonicotinoids dinotefuran, imidacloprid, and the metabolite 6-chloronicotinic acid. **(E)** Competitive binding curve showing fluorescence intensity in the presence of various natural products (p-coumaric acid, caffeine, nicotine), the carbamate insecticide carbaryl, the acaricide amitraz, and organophosphates (malathion, coumaphos, chlorpyrifos, diazinon). **(F)** Competitive binding curve showing fluorescence intensity in the presence of the pyrethroids permethrin, fenvalerate, deltamethrin, and the metabolite 3-phenoxybenzaldehyde. **(G)** Chemical structures of ligands that bind to the H-site of AmGSTD-1 based on the results of competitive fluorescence binding assay.

**Table 4.** Binding data of different ligands to AmGSTD1.

| **Ligand** | **IC50 (mM)** | **K<sub>i</sub> (μM)** |
| --- | --- | --- |
| GSH | - | - |
| CDNB | 1.22 ± 0.055 | 49.6 |
| Ethacrynic Acid | 1.02 ± 0.040 | 41.5 |
| Paraquat | - | - |
| Dicamba | - | - |
| Atrazine | - | - |
| Triclopyr | 1.86 ± 0.19 | 76.1 |
| 2,4-D | 1.01 ± 0.056 | 41.4 |
| TCP | 0.613 ± 0.027 | 25.0 |
| Chlorothalonil | - | - |
| Propiconazole | - | - |
| Myclobutanil | 3.64 ± 1.3 | 148 |
| Dinotefuran | - | - |
| 6-Chloronicotinic acid | - | - |
| Imidacloprid | 3.18 ± 0.637 | 130 |
| p-Coumaric acid | - | - |
| Caffeine | - | - |
| Nicotine | - | - |
| Carbaryl | - | - |
| Amitraz | - | - |
| Malathion | - | - |
| Coumaphos | - | - |
| Diazinon | - | - |
| Permethrin | - | - |
| Fenvalerate | - | - |
| Deltamethrin | - | - |
| 3-Phenoxybenzaldehyde | 7.40 ± 4.7 | 302 |

### 3.5. Antioxidative property of AmGSTD1

To evaluate the ability of AmGSTD1 to protect against oxidative stress, *E. coli* overexpressing AmGSTD1 was exposed to cumene hydroperoxide, hydrogen peroxide, and paraquat, which are widely known inducers of reactive oxygen species (Sanz et al., 2006; Guesmi et al., 2018). After 12 hours of exposure, the diameters (cm) of halo zones around the chemical-soaked filters in treatment (bacterial cells containing recombinant *AmGSTD1*) samples were significantly smaller than those in the control (pET-9BC) at each concentration of both cumene hydroperoxide (Fig. 7B; Fig. S1), hydrogen peroxide (Fig. 7C; Fig. S1), and paraquat treatments (Fig. 7D; Fig. S1). Compared to the control, the cumene hydroperoxide and paraquat treatments showed about 33% and 25% halo reduction, respectively (Fig. 7B; 7D). However, about 50% halo reduction was observed in hydrogen peroxide (Fig. 7C). The results obtained from disc diffusion assays indicate that AmGSTD1 exhibits the ability to protect cells against oxidative stress.

**Figure 7.**
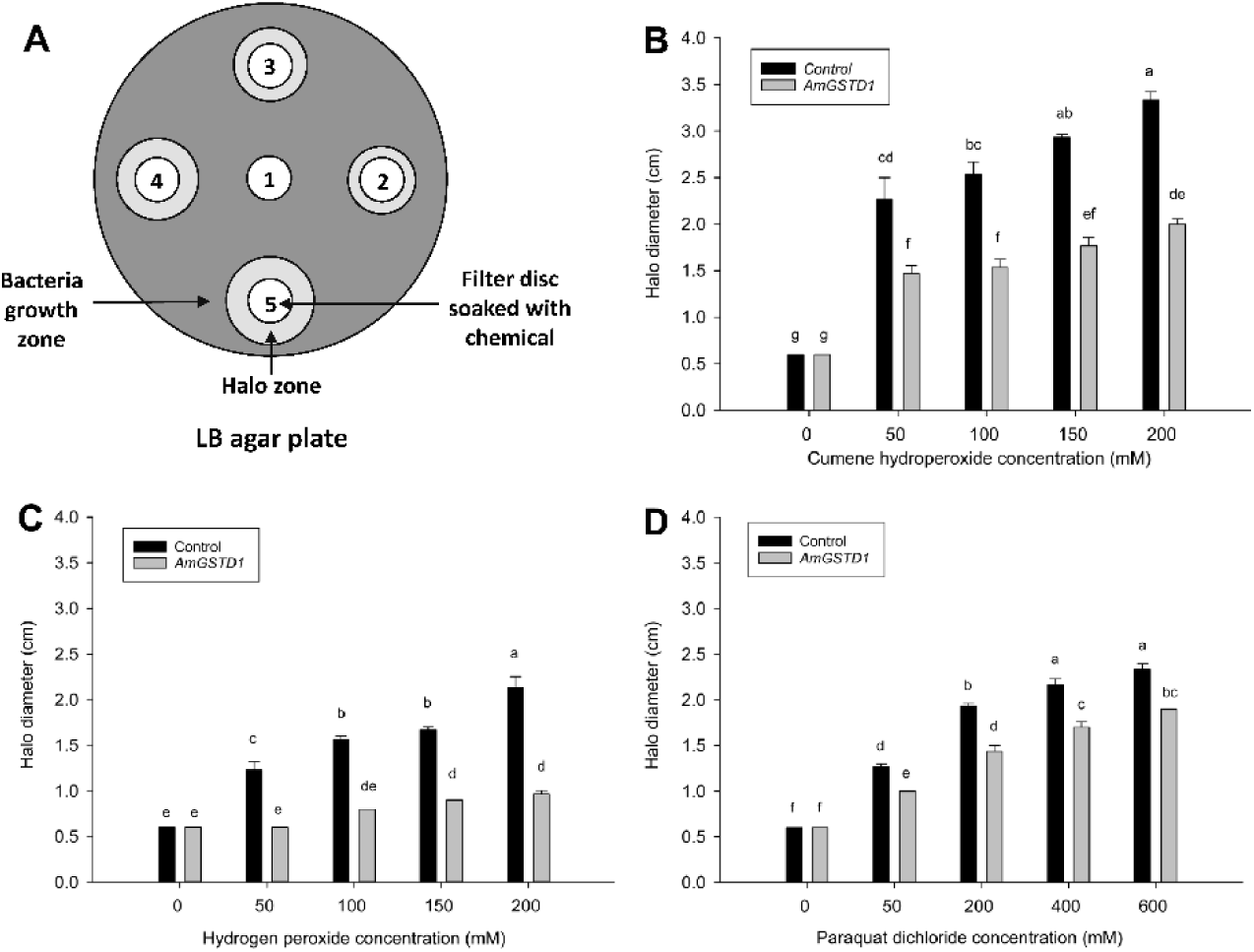
Disc diffusion assay using *E. coli* overexpressing AmGSTD1. **(A)** Cartoon illustrates the experiment design in a LB agar plate. AmGSTD1 was overexpressed in *E. coli* and approximately 5 × 10^8^ cells were plated on the plates with the antibiotics, ampicillin (200 mg/ml) and chloramphenicol (30 microgram/ml). Bacteria transfected with the pET-9BC vector only were used as the control. Bacteria transfected with pET-9BC-AmGSTD1 were used as the treatment. Filter discs were soaked with cumene hydroperoxide or hydrogen peroxide with the concentration labelled: 0 for 0 mM; 1 for 50 mM; 2 for 100 mM; 3 for 150 mM; and 4 for 200 Mm. After 12 hours of incubation at 37°C, the diameters of halo zones from the control and treatment plates were recorded and compared between the treatment and control at different concentrations of: **(B)** cumene hydroperoxide, **(C)** hydrogen peroxide, and **(D)** paraquat. Different letters indicate a significant difference in gene expression at P < 0.05, according to one-way ANOVA with the Tukey HSD test.

### 3.6. Spatial expression of AmGSTD1

The spatial expression of *AmGSTD1* in both nurse and forager bees were analyzed by qRT-PCR. Tissues such as heads, fat bodies, Malpighian tubules, midgut, legs, and muscles were used to investigate *AmGSTD1* expression. The statistical analysis shows a significant difference among the tissues for *AmGSTD1* expression (P<0.001) in both nurse and forager honey bees. The mean relative expression of *AmGSTD1* in nurse bees was highest in Malpighian tubules, followed by fat body, midgut, head, leg, and muscle (Fig. S2A). A similar expression pattern was observed in forager bees (Fig. S2B).

### 3.7. Phylogenetic relationship of AmGSTD1 with other insect GSTs

The nucleotide sequence obtained from short-read sequencing of *AmGSTD1* shows 100% identity with the GST D1 isoform X1 gene of *Ap. melifera* available in the NCBI database (Accession number: XP_006563394.1). The coding sequence of *AmGSTD1* is 720 bp long, which encodes an enzyme with a length of 240 aa. The molecular weight of AmGSTD1 is 29.42 kDa and the isoelectric point is 7.219. The phylogenetic analysis shows that AmGSTD1 was clustered into the delta clade with its predicted isomer (accession number: NP 001171499.1) and two other delta class GSTs from *T. castaneum* (accession numbers: XP 015834775.1 and XP 974273.1) with sequence similarity of 59% and 62%, respectively (Fig. 8).

**Figure 8.**
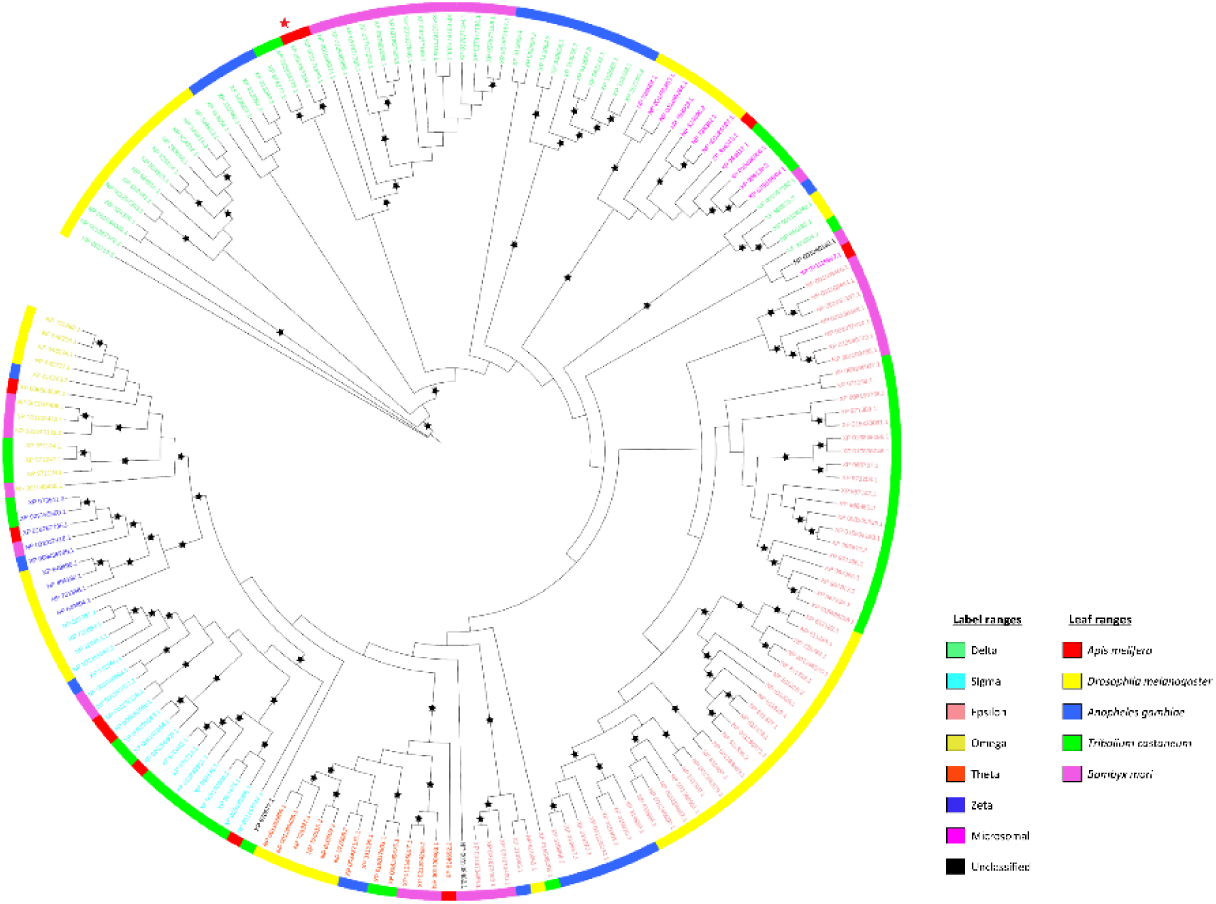
Phylogenetic analysis of AmGSTD1 with GST homologs in other insects. 182 GST amino acid sequences from five different insect species; *Apis melifera, Bombyx mori*, *Drosophila melanogaster*, *Anopheles gambiae*, and *Tribolium castaneum* were retrieved from NCBI (Table S2). Multiple sequence alignment was performed using ClustalW in MEGA X with default parameters (gap open penalty: 10, gap extension penalty: 0.2). The maximum likelihood unrooted phylogenetic tree was inferred using RaxML 8.2.12. Bootstrap analysis was performed in 500 replicates to infer the consensus tree. There are two color labels. The first color is provided to the label of the tree and different colors represent GST proteins from different classes. Next color is given to the outer leaf of the tree and color indicates insect species. Red star indicates position of AmGSTD1 in phylogenetic tree. All sequence information was listed in Table S2.

## 4. Discussion

This study presents the characterization of AmGSTD1, a delta class GST in honey bees, including its three-dimensional structure, steady-state enzyme kinetics with classic substrates, and binding data with agricultural pesticides and natural chemicals. For the first time, the 3D structure of a honey bee GST was determined with protein crystallography, providing new insights into its molecular structure. Additionally, using a fluorescence binding assay, the *K*_d_ of ANS to AmGSTD1 was found to be 2.09 ± 0.102 μM. Competitive binding assays using ANS as a fluorescence reporter in competition with various agricultural pesticides and natural products were performed to determine the capability of AmGSTD1 to bind and potentially metabolize the pesticides and natural products we assayed. It was discovered that the binding of ANS to AmGSTD1 can be inhibited with various herbicides, fungicide, insecticides, and metabolites, indicating that AmGSTD1 likely plays a crucial role in protecting honey bee health from stress imposed by agrochemicals. These findings enhance our understanding of honey bee detoxification mechanisms and their response to environmental challenges, which is vital for promoting honey bee well-being and mitigating risks associated with agrochemical exposures.

The 3D structure of AmGSTD1 and site-directed mutagenesis revealed that S34 located in the G-site is a critical catalytic residue, as the SER34 OG atom was located 3.26 Å from the complexed SG2 atom of GSH. In addition, the largest decrease in *k*_cat_ observed in the current study was for the S34A mutation, which decreased from 385 ± 11 min^-1^ in the wild-type enzyme to 92.8 ± 2.0 min^-1^ in the mutant. The catalytic serine located in the G-site is commonly found in delta class GSTs [86, 118]. Moreover, the H-site located adjacent to the G-site and completing the active site is mainly hydrophobic in character and lined with the amino acids Y130, Y138, F142, M228, and F229; the hydrophobic and aromatic character of the H-site can allow for multiple pi-pi and halogen-pi bonding interactions, pointing toward the ability of AmGSTD1 to bind and potentially metabolize a variety of hydrophobic, aromatic, and/or halogenated compounds. These structural findings are consistent with the enzyme kinetics assay results, which showed that AmGSTD1 can metabolize the model substrates CDNB, PNA, the isothiocyanates PIETC, PITC, and the oxidation byproduct HNE. GSTs have long been characterized by using multiple model substrates, natural products, and pesticides by using a combination of enzyme activity and inhibition assays [99–101, 119].

Along with structural and enzyme kinetics characterization of AmGSTD1, we performed competitive binding assays using ANS as a fluorescence reporter to assess the ability of various pesticides and natural products to bind to AmGSTD1. We made an intriguing observation that two herbicides, triclopyr and 2,4-D, as well as the triclopyr metabolite TCP, were able to inhibit fluorescence by 30-50% (Fig. 6B). Of the ligands screened, TCP was found to be the tightest binder with a *K*_i_ = 25.0 μM (Table 4). In addition to being a metabolite of the herbicide triclopyr, TCP is also the hydrolysis product of the organophosphate chlorpyriphos [120, 121]. In our study, TCP was the only compound able to displace more than 50% of ANS from AmGSTD1. Triclopyr and 2,4-D were able to displace binding of ANS, with fluorescence inhibition of 30% and 45%, with calculated *K*_i_ values of 76.1 μM and 41.4 μM, respectively, and were found to be close to the values obtained for known GST substrates CDNB and EA (Fig. 6B; Table 4). These chemicals are commonly used as herbicides in agricultural and horticultural practices, and their potential presence in the environment is a subject of concern. Although these herbicides generally exhibit low levels of toxicity, honey bees may be exposed to elevated concentrations of herbicides when they come into contact with contaminated pollen, nectar, and water sources in proximity to agricultural areas during the bloom seasons of bee-attractive crops [11]. Recent research has indicated that certain herbicides can affect honey bee health by reducing the abundance of dominant gut microbiota species and increasing the mortality of young workers when they are exposed to pathogen infections [23]. In bumblebees, it was reported that long-term exposure of glyphosate herbicide decreased collective thermoregulation by more than 25% during the period of resource limitation [122]. We also conducted additional tests on three commonly used fungicides, namely chlorothalonil, propiconazole, and myclobutanil, alongside the herbicides. Among these fungicides, only myclobutanil exhibited significant fluorescence inhibition, as shown in Fig. 6C. Previous studies have detected myclobutanil in pollens at concentrations of approximately 1 ppm [10, 70]. Myclobutanil belongs to the triazole fungicide class, which acts as a sterol biosynthesis inhibitor by targeting a cytochrome P450 (sterol 14α-demethylase, CYP51) in fungi. Exposure to myclobutanil has the potential to disrupt metabolic detoxification of plant allelochemicals in pollen and may adversely affect energy production in honey bees [123]. Recent evidence suggests that fungicides may also impact honey bee health through enhanced gut pathogen infection [66, 124]. Moreover, we found that the neonicotinoid imidacloprid and the pyrethroid insecticide metabolite 3-phenoxybenzadheyde were able to displace ANS from AmGSTD1, reducing fluorescence intensity by about 20% (Fig. 6D and 6F), suggesting they are at least weak binders to the H-site in AmGSTD1. These findings provide compelling evidence that AmGSTD1 potentially plays a critical role in safeguarding honey bee health against a broad range of agrochemicals, particularly herbicides, fungicides, and insecticides by directly metabolizing the parent compounds and/or through phase II conjugation reactions.

Previous studies have shown that AmGSTD1 is significantly up-regulated by nicotine and organophosphate (coumaphos) exposure, suggesting its potential involvement in nicotine and coumaphos detoxification processes related to these substances [80, 125]. In our competitive binding assay results (Fig. 6), we observed no significant binding for natural toxins such as p-coumaric acid, caffeine and nicotine, as well as organophosphate pesticide coumaphos and several other pesticides, indicating AmGSTD1 may not be directly involved in their metabolism. However, it is essential to consider that AmGSTD1 could still play a role in protecting honey bees from these compounds through other mechanisms, such as phase II conjugation reactions with phase I metabolites of these compounds and/or antioxidative stress (Fig. 6; Table 4).

It is well known that exposure to agrochemicals can increase oxidative stress within honey bee cells, leading to the generation of reactive oxygen species (ROS) that pose a potential threat to honey bee survival [126, 127]. GSTs, including AmGSTD1, not only participate in the conjugation of xenobiotics but also function as antioxidant enzymes, offering protection against oxidative stress induced by agrochemical exposure [61, 63]. Based on our disc diffusion assay, it was observed that AmGSTD1 displays antioxidative properties against commonly used inducers of ROS (Fig. 7). This finding aligns with a study conducted in *Apis cerana cerana*, where AccGSTD, sharing 79% amino acid sequence similarity with AmGSTD1, exhibited similar characteristics [128]. Given that AmGSTD1 is the only delta class GST in the honey bee genome, it is likely to play a critical role in honey bees’ adaptation to agrochemicals by serving as an important component in their defense against oxidative stress.

The spatial expression study for *AmGSTD1* revealed its expression in all tested tissues, with relatively higher expression levels observed in the Malpighian tubules of both nurse and forager bees (Fig. S2). Our results are consistent with a recent study by Maiwald et al., which revealed that Malpighian tubules were the predominant tissue for most analyzed detoxification genes in honey bee, including *CYP9Q2*, *CYP9Q3*, *AmGSTD1*, and *AmGSTS1* [129]. The prominence of *AmGSTD1* expression in the Malpighian tubules is of particular interest given the essential role of these organs as main excretory tissues in most insects [130]. Malpighian tubules are widely recognized as playing a key role in the production of primary urine and osmoregulation, selectively reabsorbing water, ions, and solutes, as well as excretion of natural and synthetic toxins [130, 131]. In *D. melanogaster*, dietary exposure to phenol and insecticide synergist piperonyl butoxide enhanced gene expression of multiple GSTs and increased secretion of the organic anion methotrexate in Malpighian tubules, suggesting Malpighian tubules are associated with both detoxification and excretion [131]. Several other studies have also emphasized the significant role of the Malpighian tubules in GST-mediated xenobiotic adaptation [132–134]. Therefore, considering the relatively high expression of *AmGSTD1* in Malpighian tubules of nurse and forager bees, it is plausible to propose that *AmGSTD1* serves as an important detoxification gene associated with agrochemical adaptation in honey bee.

## 5. Conclusion

Overall, our study solved the 3D structure, identified the catalytic site of a honey bee delta GST (AmGSTD1), and examined its enzymatic kinetics, substrate spectrum, along with antioxidative properties and spatial expression. The ability of AmGSTD1 to bind and interact with numerous classes of agrochemicals underscores its significance as a key detoxification enzyme in honey bees. By facilitating the binding and subsequent elimination or transformation of these harmful compounds and through defense against oxidative stress, AmGSTD1 contributes to the protection of honey bee physiology and overall health. Further investigations are required to elucidate the precise mechanism underlying the interaction between AmGSTD1 and these agrochemicals, as well as to explore its potential implications for honey bee population and pollinator conservation efforts.

## Acknowledgements

This work was supported by a faculty start-up fund from Pennsylvania State University, and the USDA National Institute of Food and Federal Appropriations under Hatch Project #PEN04770 and Accession #1010058 (F.Z.). T.W.M. was supported by USDA NIFA postdoctoral fellowship, grant #2020-67034-31780/project accession#1022959 (2020-2022). We are grateful to Avehi Singh, Kate Anton, Robyn Underwood, Margarita López-Uribe (The Pennsylvania State University) for their help in sample collection. We also thank the members of the Cornell High Energy Synchrotron and The Structural Biology Center (SBC) at the Advanced Photon Source, Argonne National Laboratory for their help with setting beamlines and data collection. We appreciate the assistance from the members at the X-ray Core facilities of Penn State University.

---

**Figure S1.**
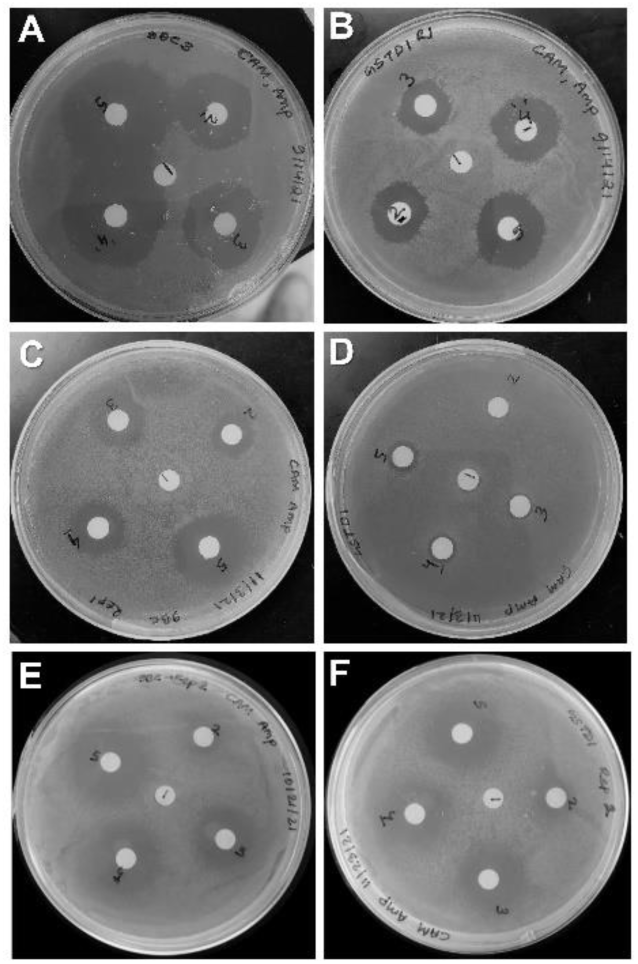
Disc diffusion assay for AmGSTD1. The bacterial discs illustrate halo diameters under stress from different concentrations of cumene hydroperoxide for the control pET-9BC vector alone **(A)** and the treatment, pET-9BC-AmGSTD1 **(B)**; the bacterial discs illustrate halo diameters under stress from different concentrations of hydrogen peroxide for the control pET-9BC vector alone **(C)** and the treatment, pET-9BC-AmGSTD1 **(D)**; and the bacterial discs illustrate halo diameters under stress from different concentrations of paraquat for the control pET-9BC vector alone **(E)** and the treatment, pET-9BC-AmGSTD1 **(F)**.

**Figure S2.**
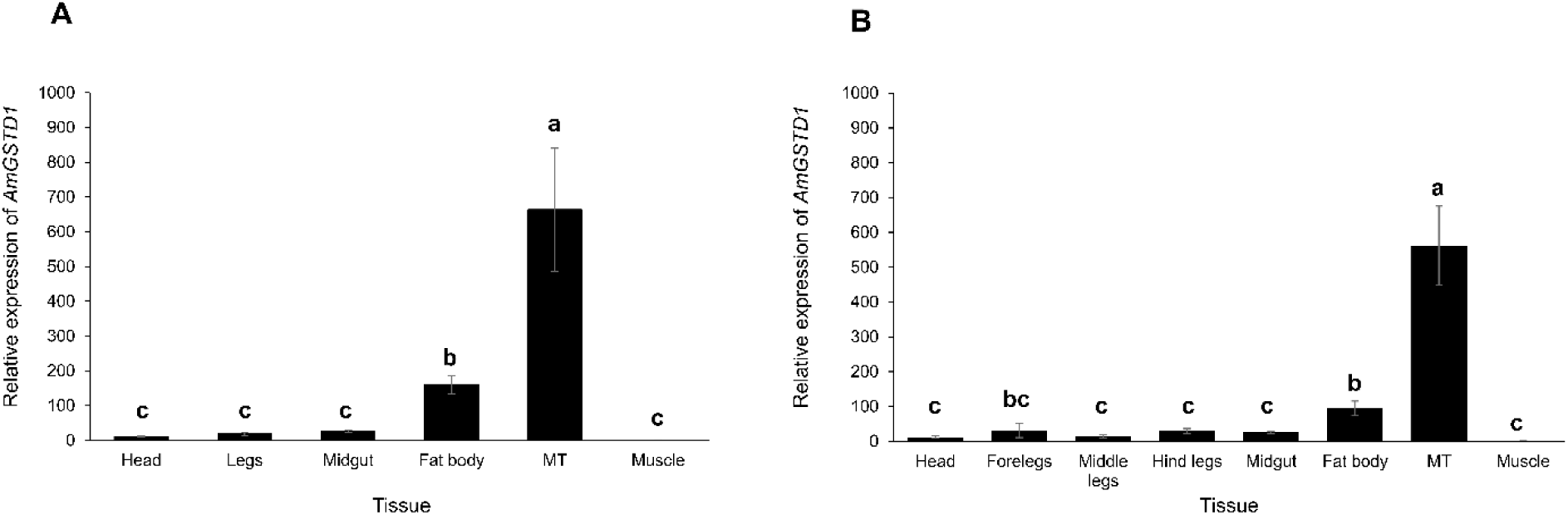
Spatial expression profile of *AmGSTD1* in nurse bees **(A)** and forager bees **(B)**. Different letters indicate a significant difference in gene expression at P < 0.05, according to one-way ANOVA with the Tukey HSD test. Note: MT-Malpighian tubules.

Table S1. Primers used for this study.

Table S2. Protein sequences of insect GST genes used to construct the maximum likelihood phylogenetic tree.

